# EmbedSimScore: Advancing Protein Similarity Analysis with Structural and Contextual Embeddings

**DOI:** 10.1101/2024.09.26.615264

**Authors:** Gourab Saha, Md Toki Tahmid, Md. Shamsuzzoha Bayzid

## Abstract

Accurately computing protein similarity is challenging due to the intricate interplay between local substructures and the global structure within protein molecules. Traditional metrics like TM-score often focus on aligning the global structures of the proteins in a rather geometry-based algorithmic way, potentially overlooking critical local-global relations and contextual comparisons. We introduce EmbedSimScore, a novel self-supervised method that generates structural and contextual embeddings by jointly considering both local substructures and global proteins’ structures. Utilizing contrastive language-structure pre-training (CLSP) and structural contrastive learning, EmbedSimScore captures comprehensive features across different scales of protein structure. These embeddings provide a more precise and holistic means of computing protein similarities, resulting in the identification of intrinsic relations among proteins that traditional approaches overlook.

## 1 Introduction

Accurately computing protein similarities is a fundamental challenge in computational biology, with significant implications for understanding protein function, evolution, and interactions. Traditional metrics such as TM-score [1] focus primarily on global structural alignments, which may overlook subtle yet critical local structural features and contextual information encoded in protein molecules. These limitations hinder the ability to fully capture the multifaceted nature of protein similarities, particularly when proteins share functional similarities despite low sequence or structural identity.

Recent advances in self-supervised learning have demonstrated the potential of deep learning models to extract meaningful representations from large datasets without explicit labels [2, 3, 4], including domains of biology[5, 6, 7]. In the context of proteins, self-supervised models have been employed to learn from amino acid sequences [8] and structures [9], but often focus on either the global structure or the sequence alone, without effectively integrating local structural nuances and contextual information.

In this work, we introduce **EmbedSimScore**, a novel self-supervised method designed to generate structural and contextual embeddings for proteins by jointly considering local substructures, global architecture, and sequence context. Our approach leverages a combination of techniques to enhance the capture of protein similarities:

- **Structural Alignment of Multiscale Subgraphs**: We adapt the self-supervised knowledge distillation framework introduced by DINO [10] for protein graphs to align representations of local and global subgraphs, ensuring that local structural nuances contribute effectively to the overall protein embedding.
- **Incorporating Contextual Similarity via Language Models**: By integrating embeddings from pre-trained protein language model ESM [5], we enrich the structural embeddings with contextual information derived from amino acid sequences, capturing functional and evolutionary relationships that may not be apparent from structure alone.
- **Contrastive Learning between Subgraphs**: We employ contrastive learning to refine local structural representations, encouraging the model to distinguish between similar and dissimilar substructures across different proteins.

## 2 Methodology

In this section, we discuss the key components of **EmbedSimScore** (Figure 1). First, we present the local and global structural alignment approach. Then, we focus on integrating contextual information through structure-language model joint contrastive learning. Finally, we explain how these components are combined into **EmbedSimScore** for comprehensive structure representation learning.

**Figure 1:**
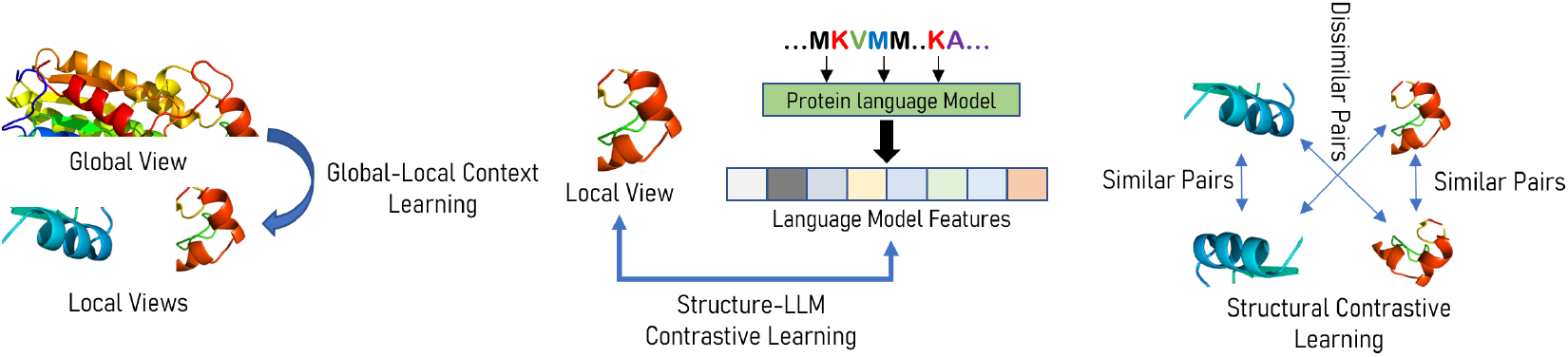
Overall training process of EmbedSimScore. Three different aspects of structural-contextual learning is employed: global-local context learning, structure-language model constrastive learning, and structural contrastive learning.

### 2.1 Structural Representation Alignment

To capture both local and global structural features, we generate multiple views of each protein graph 𝒢, inspired by the advances of DINO [10] in computer vision. We generate a **Global View** 𝒢_global_, representing the larger protein substructure, and a **Local View** 𝒢_local_, focusing on local substructures. In this setting of knowledge distillation, the **teacher network** *f*_*t*_ operates on the global view, and the **student network** *f*_*s*_ operates on both global and local views. Both networks share the same architecture but have separate parameters. The student network is trained to align its representations with those of the teacher network by minimizing the alignment loss 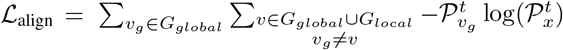, where 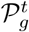 and 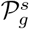 are the probability distribution of the pseudo-labels for graph *g* produced by *f*_*t*_ and *f*_*s*_ respectively.

### 2.2 Integrating Contextual Similarity via Language Models

To incorporate contextual information from protein sequences, we utilize a pre-trained language model ℳ that generates sequence embeddings **h**_*S*_ from the amino acid sequence *S* corresponding to the protein graph 𝒢. The structural embeddings **h**_𝒢_ of the student network are aligned with the sequence embeddings using a contrastive loss inspired by CLIP [11]. The loss is defined as 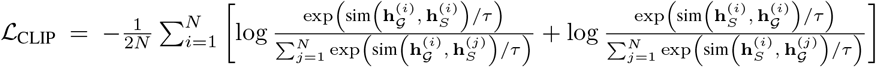, where sim (·, ·) denotes the cosine similarity, and *τ* is a temperature parameter controlling the sharpness of the distribution.

### 2.3 Contrastive Learning Between Subgraphs

To improve local structural representation learning, we apply contrastive learning between the augmented views of local subgraphs. For each protein, we create two augmented versions of a local subgraph, resulting in embeddings **h**^(*i*,1)^ and **h**^(*i*,2)^. The contrastive loss is defined as 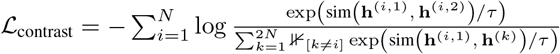.

### 2.4 Overall Objective

The total loss function combines the above components: ℒ = ℒ_align_ + *λ*_context_ ℒ_context_ + *λ*_contrast_ ℒ_contrast_, where *λ*_align_, *λ*_context_, and *λ*_contrast_ are hyperparameters that balance the contributions of each loss term. With this combined approach, we train a graph neural network (GNN) backbone with five layers of GVP [12] on 48k proteins curated from protein data bank(PDB) [13] and their respective language model features generated using ESM-2[5] 650M model for 300 epochs. This backbone can embed any protein sequence, enabling the computation of structural and contextual relationships between proteins by comparing their **EmbedSimScore** embeddings.

## 3 Results

In this section, we discuss our findings on proteins’ structural and contextual similarity calculation on a representative set with 50 protein molecules for whom we extract their 3d structure from the corresponding PDB file. This selection of protein structures is diverse across different protein families.

In Figure 2 (a), we plot the TM-score between each pair of protein molecules in our representative set. A score less than 0.2 has been discarded as means that the similarities are rather random [14]. In Figure 2 (b), we have plotted the embedding similarity between each pair of protein embedding generated by EmbedSimScore. From the figure, we see that 24 proteins pairs that are identified as similar by TM-score metrics are all identified by EmbedSimScore embedding as similar. However, EmbedSimScore captures another 7 proteins which are structurally and contextually similar as calculated from their embedding similarity (similarity score > 0.98).

**Figure 2:**
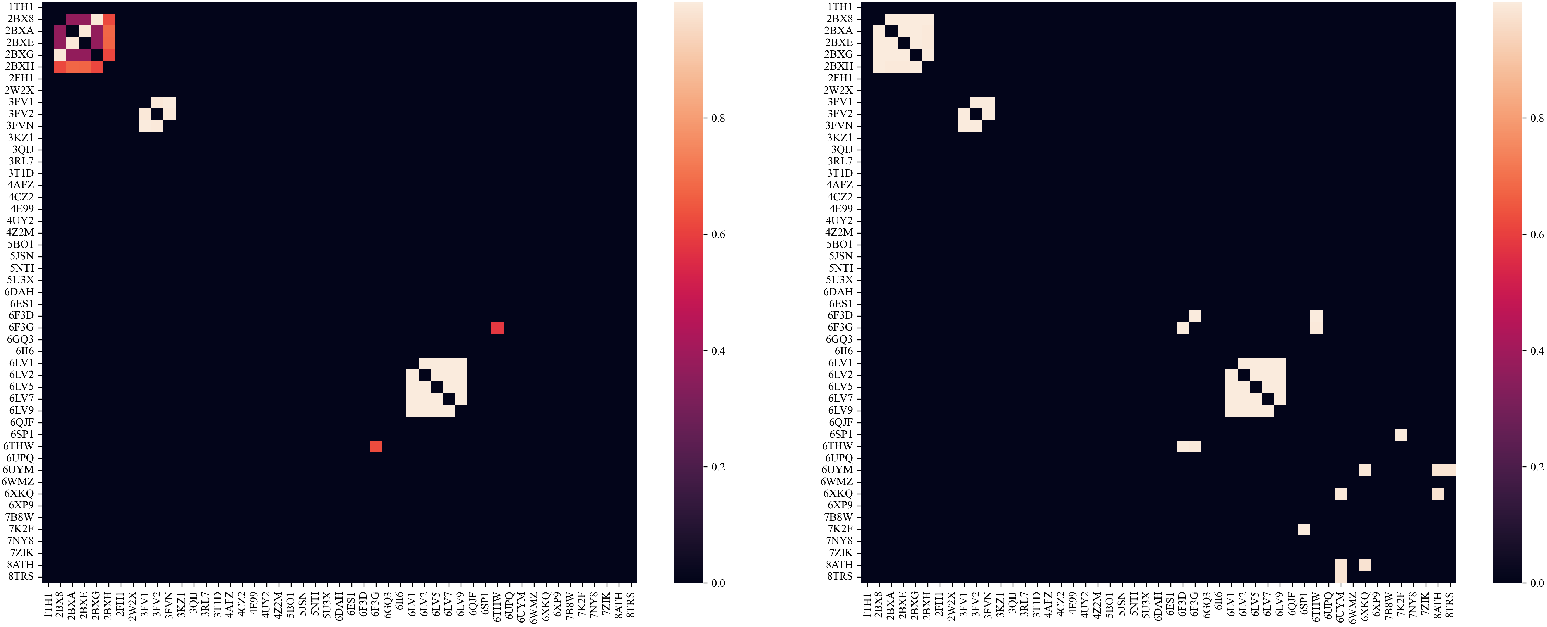
The heatmap in figure (a) represents similarity as shown by TM-score, and the one in figure (b) represents similarity scores as predicted by EmbedSimScore

In Table 3, we show the 3D structural configurations of each pair of proteins that are identified as similar by EmbedSimScore, but not captured with TM-score. We see that, most of them have local structural similarity (similar folding structure in subgraphs), even though the overall global conformations may vary. Note that, we did not compare the structural configuration of 6F3D-6F3G pair, as they are the different conformation of the same protein molecule.

**Table 1:**
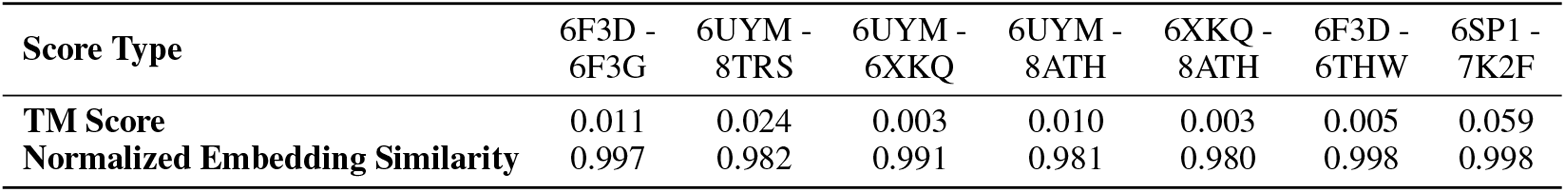
Comparison of TM and Embedding Similarity Scores for 3D Structure of PDB Pairs.

**Table 2:**
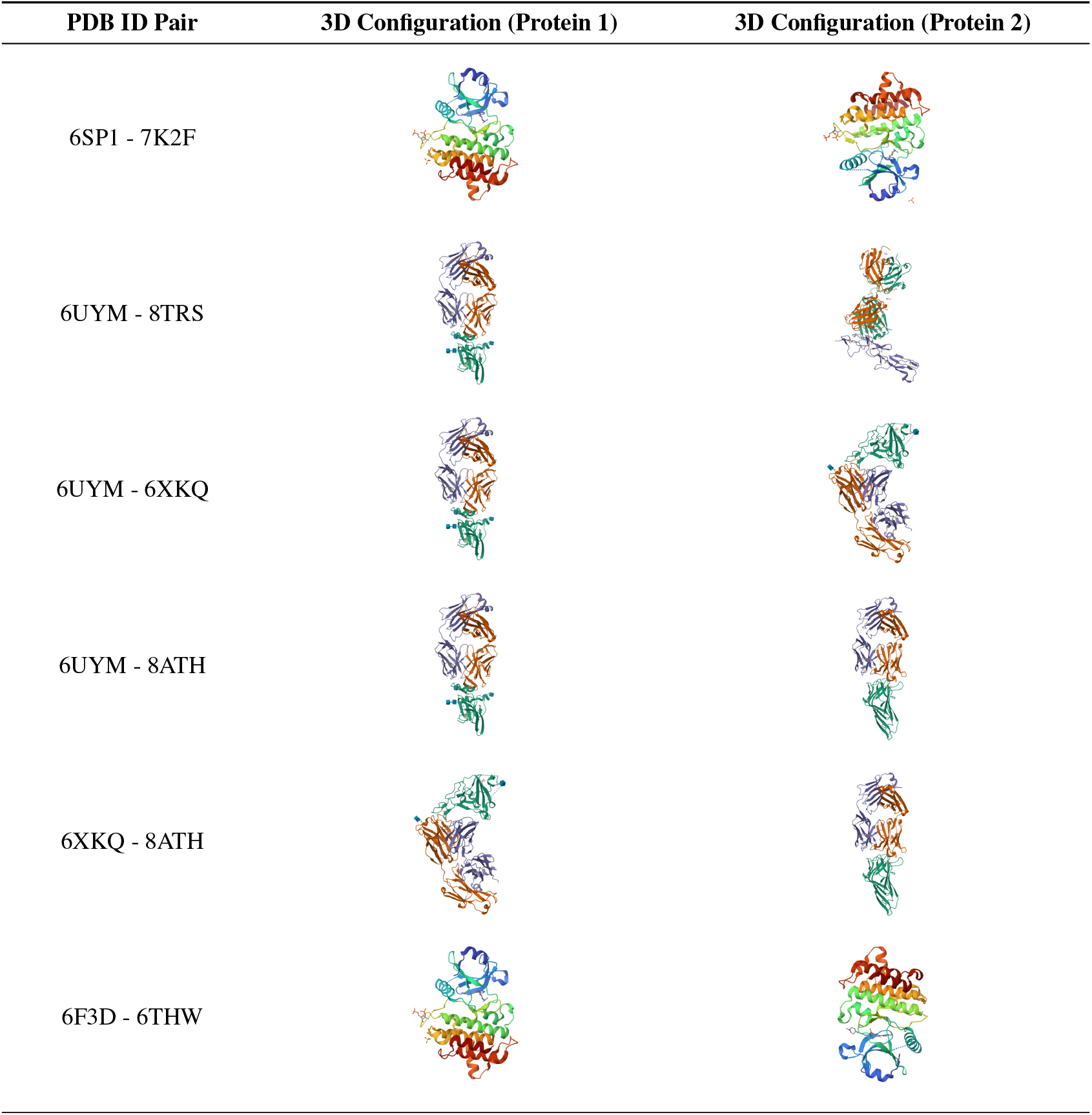
PDB pairs and their respective conformations.

## 4 Conclusion

We presented **EmbedSimScore**, a self-supervised approach that enhances protein similarity computation by considering both local substructures and global protein features. Using structural alignment, contrastive learning, and language model integration, **EmbedSimScore** generates embeddings that capture relationships between proteins, often overlooked by traditional metrics like TM-score. **EmbedSimScore** sets a promising direction for advancing protein similarity analysis. Notably, aligning two proteins through their local substructure opens up new possibilities for various tasks, including protein engineering with specific functions (e.g., designing enzymes that catalyze novel reactions), drug discovery (e.g., identifying proteins with similar binding pockets for new therapeutic targets), and more, offering a powerful tool for targeted applications in these fields.

## References

[1] Yang Zhang and Jeffrey Skolnick. Scoring function for automated assessment of protein structure template quality. Proteins: Structure, Function, and Bioinformatics, 57(4):702–710, 2004.

[2] Jacob Devlin. Bert: Pre-training of deep bidirectional transformers for language understanding. arXiv preprint 1810.04805, 2018.

[3] Kaiming He, Xinlei Chen, Saining Xie, Yanghao Li, Piotr Dollár, and Ross Girshick. Masked autoencoders are scalable vision learners. 2111.06377, 2021.

[4] Kaiming He, Haoqi Fan, Yuxin Wu, Saining Xie, and Ross Girshick. Momentum contrast for unsupervised visual representation learning. In Proceedings of the IEEE/CVF conference on computer vision and pattern recognition, pages 9729–9738, 2020.

[5] Zeming Lin, Halil Akin, Roshan Rao, Brian Hie, Zhongkai Zhu, Wenting Lu, Allan dos Santos Costa, Maryam Fazel-Zarandi, Tom Sercu, Sal Candido, et al. Language models of protein sequences at the scale of evolution enable accurate structure prediction. BioRxiv, 2022:500902, 2022.

[6] Walid Ahmad, Elana Simon, Seyone Chithrananda, Gabriel Grand, and Bharath Ramsundar. Chemberta-2: Towards chemical foundation models. arXiv preprint 2209.01712, 2022.

[7] Zhihan Zhou, Yanrong Ji, Weijian Li, Pratik Dutta, Ramana Davuluri, and Han Liu. Dnabert-2: Efficient foundation model and benchmark for multi-species genome. arXiv preprint 2306.15006, 2023.

[8] Roshan Rao, Nicholas Bhattacharya, Neil Thomas, Yan Duan, Peter Chen, John Canny, Pieter Abbeel, and Yun Song. Evaluating protein transfer learning with tape. Advances in neural information processing systems, 32, 2019.

[9] Pablo Gainza, Freyr Sverrisson, Frederico Monti, Emanuele Rodola, Davide Boscaini, Michael M Bronstein, and Bruno E Correia. Deciphering interaction fingerprints from protein molecular surfaces using geometric deep learning. Nature Methods, 17(2):184–192, 2020.

[10] Mathilde Caron, Hugo Touvron, Ishan Misra, Hervé Jégou, Julien Mairal, Piotr Bojanowski, and Armand Joulin. Emerging properties in self-supervised vision transformers. In Proceedings of the IEEE/CVF international conference on computer vision, pages 9650–9660, 2021.

[11] Alec Radford, Jong Wook Kim, Chris Hallacy, Aditya Ramesh, Gabriel Goh, Sandhini Agarwal, Girish Sastry, Amanda Askell, Pamela Mishkin, Jack Clark, et al. Learning transferable visual models from natural language supervision. In International conference on machine learning, pages 8748–8763. PMLR, 2021.

[12] Bowen Jing, Stephan Eismann, Patricia Suriana, Raphael John Lamarre Townshend, and Ron Dror. Learning from protein structure with geometric vector perceptrons. In International Conference on Learning Representations, 2020.

[13] Helen M Berman, John Westbrook, Zukang Feng, Gary Gilliland, Talapady N Bhat, Helge Weissig, Ilya N Shindyalov, and Philip E Bourne. The protein data bank. Nucleic acids research, 28(1):235–242, 2000.

[14] Jinrui Xu and Yang Zhang. How significant is a protein structure similarity with tm-score= 0.5? Bioinformatics, 26(7):889–895, 2010.

